# Discriminability of numerosity-evoked fMRI activity patterns in human intra-parietal cortex reflects behavioral numerical acuity

**DOI:** 10.1101/198002

**Authors:** G. Lasne, M. Piazza, S. Dehaene, A. Kleinschmidt, E. Eger

## Abstract

Areas of the primate intraparietal cortex have been identified as an important substrate of numerical cognition. In human fMRI studies, activity patterns in these and other areas have allowed researchers to read out the numerosity a subject is viewing, but the relation of such decodable information with behavioral numerical proficiency remains unknown.

Here, we estimated the precision of behavioral numerosity discrimination (internal Weber fraction) in twelve adult subjects based on psychophysical testing in a delayed numerosity comparison task outside the scanner. FMRI data were then recorded during a similar task, to obtain the accuracy with which the same sample numerosities could be read out from evoked brain activity patterns, as a measure of the precision of the neuronal representation. Sample numerosities were decodable in both early visual and intra-parietal cortex with approximately equal accuracy on average. In parietal cortex, smaller numerosities were better discriminated than larger numerosities of the same ratio, paralleling smaller behavioral Weber fractions for smaller numerosities. Furthermore, in parietal but not early visual cortex, fMRI decoding performance was correlated with behavioral number discrimination acuity across subjects (subjects with a more precise behavioral Weber fraction measured prior to scanning showed greater discriminability of fMRI activity patterns in intraparietal cortex, and more specifically, the right LIP region).

These results suggest a crucial role for intra-parietal cortex in supporting a numerical representation which is explicitly read out for numerical decisions and behavior.

## Introduction

Humans share with other animals the ability to rapidly extract approximate numerosity from a visual scene, and to compare numerosities with an accuracy that roughly depends on their numerical ratio (Feigenson et al., 2004; Cantlon and Brannon, 2006). Numerosity perception can be psychophysically dissociated from other quantitative judgements (Anobile et al., 2016a; Cicchini et al., 2016), suggesting it relies on dedicated neural extraction channels. In accord with this, numerosity responsive units supporting ratio-dependent discrimination can develop through unsupervised learning in hierarchical generative networks (Stoianov and Zorzi, 2012). It is remarkable that the individual precision of basic non-verbal number discrimination can be predictive of current and future higher-level symbolic arithmetic skills (Gilmore et al., 2007; Halberda et al., 2008; Anobile et al., 2016b), even though the human species’ particularly highly developed mathematical abilities undoubtedly rely on multiple foundational capacities (Butterworth, 2010; De Smedt et al., 2013).

Evidence from several neuroscientific techniques has outlined a set of brain areas with particular importance for numerical processing. In macaque monkeys, single neurons responding differentially to different numbers of perceived items have been described in sub-regions of intra-parietal and prefrontal cortex (Nieder and Miller, 2004; Roitman et al., 2007). At a coarse spatial scale, functional MRI has demonstrated increased activation during a variety of numerical as opposed to non-numerical tasks (see Arsalidou and Taylor, 2011, for a meta-analysis), and responsiveness to numerical deviance during passive viewing (e.g., Piazza et al., 2004; Cantlon and Brannon, 2006; Jacob and Nieder, 2009; He et al., 2015), in similarly located regions. In recent years, multivariate decoding methods have been introduced to make fine-grained (e.g., within the same category) discriminations between perceptual features on the basis of fMRI activity profiles across voxels (e.g., Norman et al., 2006; Tong and Pratte,2012). Using this approach it has been possible to read out the number seen or held in mind by a subject from fMRI activity in parietal areas functionally equivalent to those carrying numerical responses in macaques (Eger et al., 2009; Eger et al., 2015).

While the neuronal substrates underpinning numerosity perception have been described in some depth, it remains insufficiently understood what brain mechanisms give rise to variations in numerical performance, either across subjects or different experimental situations. The ability to read out numerosity information from a given brain area does not necessarily imply that subjects are relying on this information when making numerical discriminations. Recent fMRI studies have shown that numerosity could also be decoded from areas beyond those considered the core substrate of numerical processing, for example early and ventral visual cortex (Bulthé et al., 2014; Bulthé et al., 2015; Eger et al., 2015), though not in all cases (Castaldi et al., 2016). Controlling non-numerical properties when working with non-symbolic numerical stimuli is a complex task and often not exhaustively achieved within a single experimental context (Gebuis and Reynvoet, 2012), making positive findings in early visual areas difficult to interpret. Nevertheless, a recent EEG study using an experimental design that allowed testing the effect of variation along multiple non-numerical quantitative dimensions, observed that already very early components of the ERP, compatible with sources in early visual cortex, were modulated more by change in the numerical rather than other dimensions (Park et al., 2016). Such effects could be related to the segmentation of individual items, or the operation of a combination of spatial filters which has been proposed as a potential mechanism to extract an estimate of numerosity (Dakin et al., 2011), plausibly located at earlier levels of the visual hierarchy. Given such potential contributions of earlier visual regions to numerosity extraction, the question arises to what extent numerical acuity is limited by efficiency of the processes at these earlier, or higher-level processing stages as parietal cortex.

To shed light on the question of which, among several areas where numerosity could previously be successfully decoded, are the most critical to determine the precision of behavioral discrimination of this feature, here we related psychophysical measurements and multivariate decoding analysis of fMRI patterns, focusing on variability between subjects and across different numerical ranges. Each subject’s precision of behavioral discrimination (internal Weber fraction) was estimated based on psychophysical testing in a delayed numerosity comparison task outside the scanner. FMRI data were then recorded during a similar task, to obtain the accuracy with which the same sample numerosities could be read out from evoked brain activity patterns, as a measure of the precision of the neuronal representation.

## Materials and Methods

### Subjects and MRI acquisition

Twelve healthy volunteers (6 male and 6 female; mean age 22 years) participated in the study. All but one subject were right-handed and all had normal or corrected-to-normal visual acuity. The study was approved by the regional ethical committee (Hôpital de Bicêtre, France). Functional images were acquired on a 3 Tesla MR system with 12-channel head coil as T2*-weighted echo-planar image (EPI) volumes with 2 mm isotropic voxels. Thirty oblique transverse slices covering essentially the dorsal visual pathway and superior parts of frontal cortex were obtained in ascending interleaved order (repetition time [TR] = 2.52 s; echo time [TE] = 33ms; field of view [FOV] = 192 mm; flip angle [FA] = 84°)

### Experimental Design and Statistical Analysis

#### Stimuli and Procedure

During fMRI and in the separate behavioral experiments preceding it, subject performed a delayed numerosity comparison task on displays consisting of different numbers of randomly positioned light grey dots on a black background, inside an implicit circular area subtending ~8° of visual angle at the center of the screen (Figure 1A). Two separate stimulus sets equated either the overall area of grey (resulting in decreasing dot size with increasing number), or the individual dot size between numerosities (resulting in increasing number of grey pixels with increasing number).

**Figure 1:**
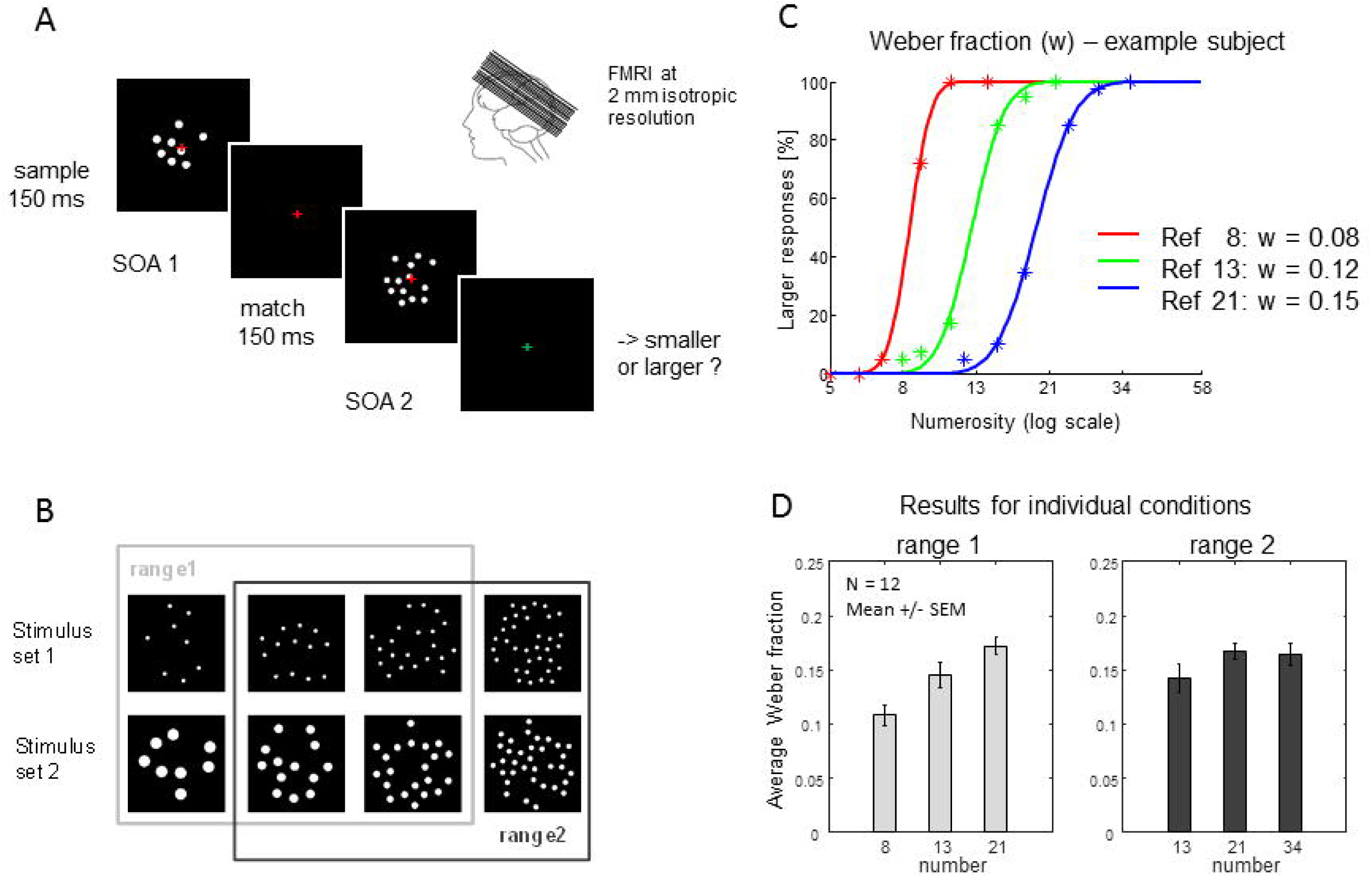
Paradigm: A) In a delayed numerical comparison task, subjects were presented with a sample numerosity for 150 ms that after an SOA of several seconds was followed by a match numerosity presented for the same short duration and required a smaller/larger judgment. The critical fMRI data used for multivariate decoding were the responses evoked by the sample numerosities. B) In each half of the experimental runs, three sample numerosities were used, forming two partly overlapping ranges (either 8, 13 and 21, or 13, 21 and 34 items). Behavioral results: The percentage of larger responses in the preceding independent behavioral experiment was fitted with cumulative Gaussian functions to obtain a measure of the internal Weber fraction (w). Panel C) illustrates the functions fitted in one subject, with steeper lopes for smaller numerosities. Panel D) displays group results (n=12, means and SEM) for w obtained for the three sample numerosities in the two ranges.

Each trial started with a sample stimulus display for 150 ms, and after a first delay, a match stimulus appeared for 150 ms; after a second delay, a new trial was presented. During the duration of a trial, a red cross was present in the center of the screen, which turned to green for 2 s after the disappearance of the match stimulus. Subjects had to memorize the approximate number of dots of the sample stimulus on each given trial and to respond by button press, after presentation of the match stimulus, depending on whether they judged the match number smaller or larger than the previous sample number.

Four different sample numerosities separated by a ratio of ~1.6 (8, 13, 21, and 34), were used. For reasons not relevant to the aim of the present report, different ranges of three sample numerosities were presented in different experimental runs (Figure 1 B), the first range comprising numerosities 8, 13, and 21, and the second range numerosities 13, 21 and 34. In the independent behavioral experiment, six match numerosities were used per sample: (sample 8: 5, 6, 7, 9, 11, and 15 items, sample 13: 8, 9, 11, 15, 18, and 22 items, sample 21: 12, 15, 18, 24, 29, and 36 items, sample 34: 20, 24, 28, 40, 48, and 58 items). Each sample display was separated from the match display by a stimulus onset asynchrony (SOA 1) of either 3 or 6s, whereas the delay between the match and the new sample presentation (SOA 2)was fixed at 3s. A behavioral run contained 72 trials drawn from the same range of sample numerosities (3 sample numerosities x 2 SOA 1 x 6 match numerosities x 2 set types controlling for different physical variables). Each subject performed a first block of 5 runs with one of the two ranges (e.g., [8-21]), and after a short pause, a second block of 5 runs with the other remaining range (e.g., [13-34]). The order of presentation of these two blocks was counterbalanced across subjects. To obtain a sufficient amount of data, two behavioral testing sessions per subject were performed under the same conditions within an interval of around two weeks.

In the fMRI experiment, to equate subjective task difficulty, match numbers were chosen based on the individual psychometric function of each subject computed from the results of the prior behavioral experiment, using psignifit toolbox (http://psignifit.sourceforge.net/) (Wichmann and Hill, 2001). Standard match stimuli were chosen to correspond to those numerosities that yielded 25/75% of larger responses in the previous behavioral testing (“standard” trials). In addition, on a small percentage of trials, match numerosities corresponded to 5/95% of larger responses (“catch” trials). The order of trials was pseudo-randomized, as well as the SOA 1 and SOA 2, which could be 3, 4, 5 or 6 s. A run contained 60 “standard” trials (3 samples x 2 matches x 2 sets x 5 repetitions) and 12 “catch” trials (3 samples x 2 matches x 2 sets), from the same range of sample numerosities. East subject performed four runs: run 1 and 4 corresponding to one of the two numerosity ranges, and runs 2 and 3 to the other range, with the order of range presentation counterbalanced across subjects. Before each run, subjects performed a short six-trial-training in which sample and (their respective) match numerosities of the following run were presented. An experiment lasted 45 minutes.

## Data analysis

To obtain measures of the internal Weber fraction of numerosity representation, the percentage of larger responses to match numbers in the psychophysical experiment as a function of the logarithmic difference between sample and match numerosities was fitted with a cumulative Gaussian function. The standard deviation of this function, when divided by √2, yields the internal Weber fraction (Dehaene, 2007).

From the Weber fractions for individual numerosities within each range, we further computed the sensitivity index (d’) for discrimination between all possible pairs of sample numerosities n1 and n2 as in the following way, where w1 is the internal Weber fraction for n1 and w2 the internal Weber fraction for n2.

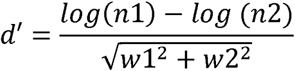

The fMRI data were preprocessed with SPM8 (http://www.fil.ion.ucl.ac.uk/spm/software), including realignment to the first volume as reference, and normalization to the standard template of the Montreal Neurological Institute (MNI) using SPM’s segment algorithm.

The normalized, unsmoothed EPI volumes were entered into a GLM, where each sample stimulus event was modeled as a condition, resulting in 72 conditions per session (60 events for “standard” trials and 12 events for “catch” trials). Match stimulus time points of the same relative magnitude (within the respective range) and response type (smaller or larger than the sample stimulus) were grouped and modeled as further trial types, resulting in 6 additional conditions (3 magnitudes x 2 response types) per session. The onsets of these 78 conditions were convolved with a standard hemodynamic response function. The resulting 60 beta estimates (per session) for the “standard” sample stimulus conditions were used in the multi-voxel pattern analysis. We defined regions of interest (ROIs) within large anatomical masks of both parietal (left and right superior and inferior parietal lobules) and early visual (left and right area 17) cortex created using WFU PickAtlas (http://fmri.wfubmc.edu/-software/pickatlas) (Maldjian et al., 2003) by selecting for each subject the 600 most significantly activated voxels in the t-contrast of all “standard” sample stimulus conditions vs baseline. In addition, the ROIs originally introduced by a previous study (Eger et al., 2015) to operationally define human equivalents of areas LIP and VIP were reapplied to the current data set, selecting for each subject within each of the 4 masks the 150 most activated voxels in the same contrast as mentioned previously.

For pattern classification, conditions were labeled according to sample numerosity, collapsing across the two different stimulus sets. Pattern recognition analysis was performed on mean-corrected trial-wise parameter estimate vectors (40 vectors / condition) using linear Support Vector Machines (SVM) with regularization parameter C = 1 in scikit-learn (http://scikitlearn.org/stable/) (Pedregosa et al., 2011). The classification was performed for all possible pairs of numerosities within each range (theoretical chance level = 50 %). For each comparison, for trial n=1:40, the n-th trial of each condition was left out for test when training the classifier, and the percentage of correct identification on left-out data was computed across the entire cross-validation cycle.

Confusion matrices were constructed from the output of classifiers from all pairwise comparisons. In addition, in analogy to the analysis of the behavioral data, the sensitivity index (d’) was computed for pairwise comparisons between numerosities as:

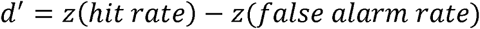

Across subject statistics on behavioral Weber fractions and fMRI classification performance used paired two-tailed t-tests, and the across subject relation between these measures was probed by Pearson correlations.

## Results

### Behavioral results

We fitted the percentage of larger responses to match numbers with cumulative Gaussian functions to obtain the internal Weber fraction (w) of the numerosity representation (see Materials and Methods for details). Fits were calculated per magnitude, range, sample match delay, and testing session, and subsequently averaged for each magnitude within each range ^1^. Figure 1C illustrates the fitted curves for one subject, and Figure 1D the estimated w across subjects for the different conditions. Weber fractions were not completely equal across numerosities in this study, but gradually increased from number 8 to 13 to 21 after which they remained constant. Pairwise statistical comparisons between conditions across the 12 subjects confirmed that Weber fractions for numerosity 8 were significantly different from the ones for all the other conditions (compared to 13 range 1: t(11) = 3.57, p = 0.00439, 21 range 1: t(11) = 5.71, p = 0.00014, 13 range 2: t(11) = 3.41, p = 0.00587, 21 range 2: t(11) = 7.72, p = 0.00001, 34: t(11) = 3.40, p = 0.00592, all paired two-tailed t-tests), while no other comparisons reached significance.

For correlation with the fMRI decoding results, an average Weber fraction across numerosities and ranges was computed for each subject. Across the group these values had a mean of 0.15 (with the minimum being 0.13, and the maximum 0.19).

### FMRI decoding results in early visual and parietal cortex

Multivariate classifiers based on support vector machines were used to discriminate between all pairs of sample numerosities within a given range, within regions of interest in early visual and parietal cortex. The detailed pattern of classification performance is displayed in Figure 2A in form of the confusion matrices obtained from the pairwise classification for both regions. Percentages of classification are plotted for the true conditions against the predicted conditions, values on the diagonal reflect correct identifications, and off-diagonal values mis-classifications. Overall accuracies and patterns of performance are rather similar for the two regions of interest, with a slight apparent difference in the accuracy across the two numerical ranges between regions: early visual cortex activation patterns allowed to identify sample numerosities with approximately equal accuracy within both ranges, while in parietal cortex classification was on average better in range 1 which included smaller sample numerosities than in range 2 (as indicated by lower diagonal and higher off-diagonal values). This pattern resembles the one observed in the behavioral results (smaller Weber fractions for smaller numerosities).

**Figure 2:**
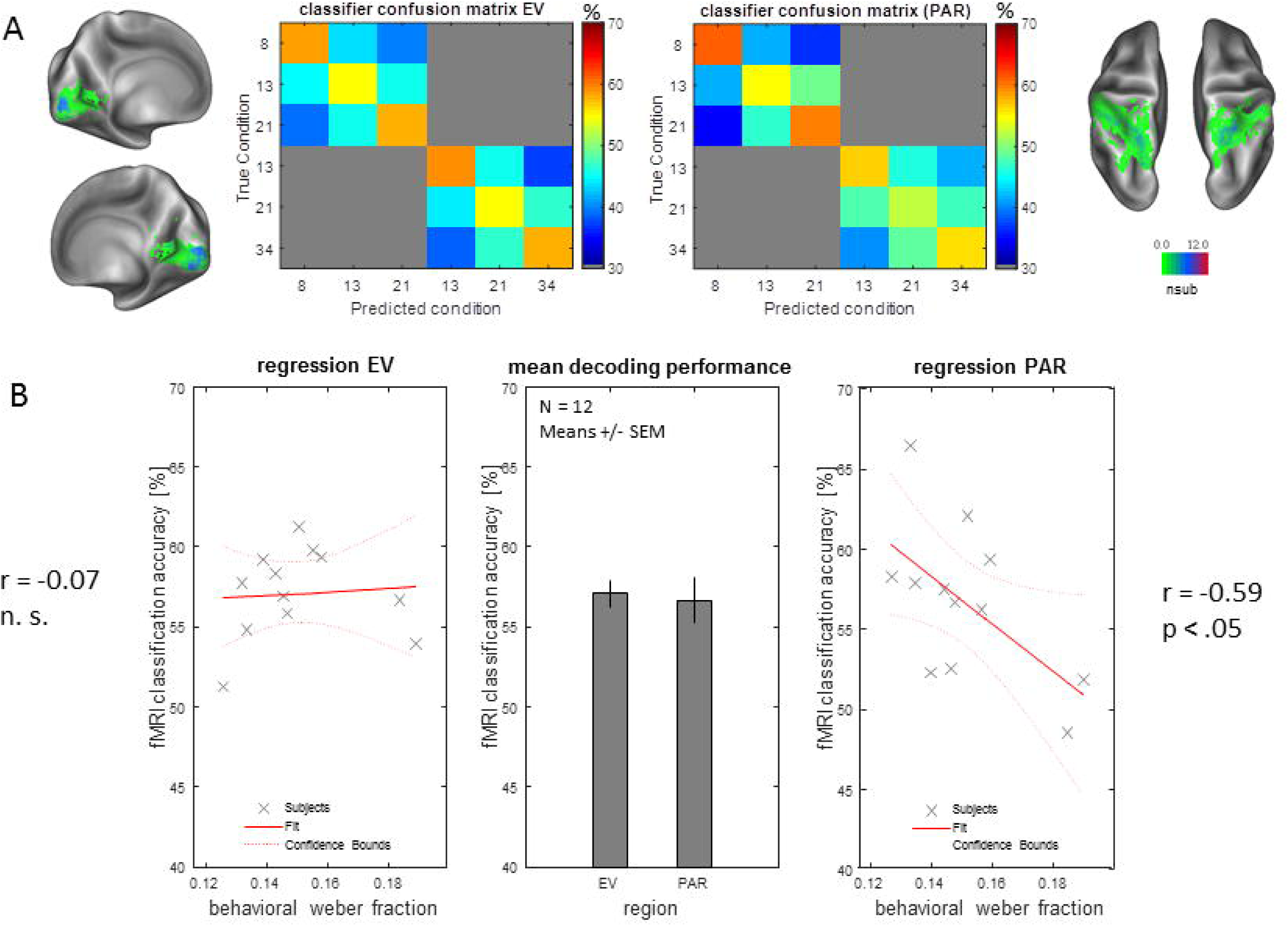
FMRI decoding results and across-subject correlation with behavioral Weber fractions: Panel A) displays the average confusion matrices obtained from all pairwise comparisons between numerosities in each numerical range, for regions of interest in early visual and parietal cortex. Individual subjects’ ROIs were defined as the 600 voxels most activated across all sample numerosities vs baseline, within each anatomical volume of interest. The color code of the ROI visualization reflects the number of subjects in whom one given voxel location was selected. In the confusion matrices, the values on the diagonal correspond to the percentage of correct identifications for each numerosity on average across all pairwise comparisons, and the off-diagonal values correspond to the percentages of incorrect identification (confusionwith one of the other numerosities). The theoretical chance level for correct identification is 50 %. Panel B) displays the average classification performance (n=12, means and SEM) for pairwise discrimination between numerosities for each ROI (middle panel) together with correlations between individual subjects’ fMRI classification performance and behavioral Weber fractions (left panel for early visual cortex, right panel for parietal cortex). In spite of a very similar average decoding performance in the two ROIs, a negative correlation between behavioral w and fMRI decoding performance was found only in the parietal ROI, indicating that subjects with a more precise behavioral discrimination measured prior to scanning had more precisely discriminable evoked activation patterns in that region.

Across all pairwise comparisons, decoding performance was on average 57.1% correct in early visual cortex and 56.7 % correct in parietal cortex (in both cases significantly different from the theoretical chance level of 50 %, paired two-tailed t(11) = 8.72, p = 0.000003 in early visual cortex, and t(11) = 4.71, p = 0.000641 in parietal cortex).

### Across subject correlation between behavioral precision and fMRI decoding performance

To test for the behavioral relevance of these occipital and parietal number representations, we capitalized on the inter-individual differences in the precision of the numerical representation as determined by psychophysics, and correlated the psychophysical Weber fractions across subjects with the average decoding accuracies. The idea underlying this approach is that if a neural representation underpins (or determines) behavior then decoding accuracy should predict psychophysical performance. Figure 2B shows the extent to which fMRI classification performance is predicted by Weber fractions from the independent behavioral experiment in both regions of interest. In spite of the very similar level of prediction accuracy on average (middle panel), a significant Pearson’s correlation of behavioral Weber fractions with fMRI decoding accuracies across subjects was only found in parietal (r = -0.59, p = 0.0451) but not early visual cortex (r =-0.07, p = 0.8305). A negative correlation in this case indicates that subjects with a higher behavioral acuity (as evidenced by smaller w) have a more precise neuronal representation (as evidenced by higher decoding performance for individual numbers from multi-voxel activation patterns).

To further explore which precise sub-regions of parietal cortex might most account for the observed correlation between behavioral performance and fMRI decoding accuracy, we probed the same relation within four parietal sub-regions that were defined in a previous study using localizer scans motivated by neurophysiology: the putative human equivalents of lateral (LIP) and ventral (VIP) intra-parietal areas of the macaque (Figure 3) in the left and right hemispheres. All these regions showed a tendency towards the expected negative correlation, as the one observed in the global parietal ROI. However, this correlation had the highest value in the right LIP region and it reached significance only in this region when testing each ROI in isolation (right LIP: r = -0.74, p = 0.0059, left LIP: r = -0.49, p = 0.1067, right VIP: r = -0.38, p = 0.2266, left VIP: r = -0.12, p = 0.7053). The correlation in the right LIP region remains significant even when applying Bonferroni correction for multiple comparisons in this case (4 subregions tested: α_altered_ =.05/4 = .0125, pcorrected = .0236 for right LIP region).

**Figure 3:**
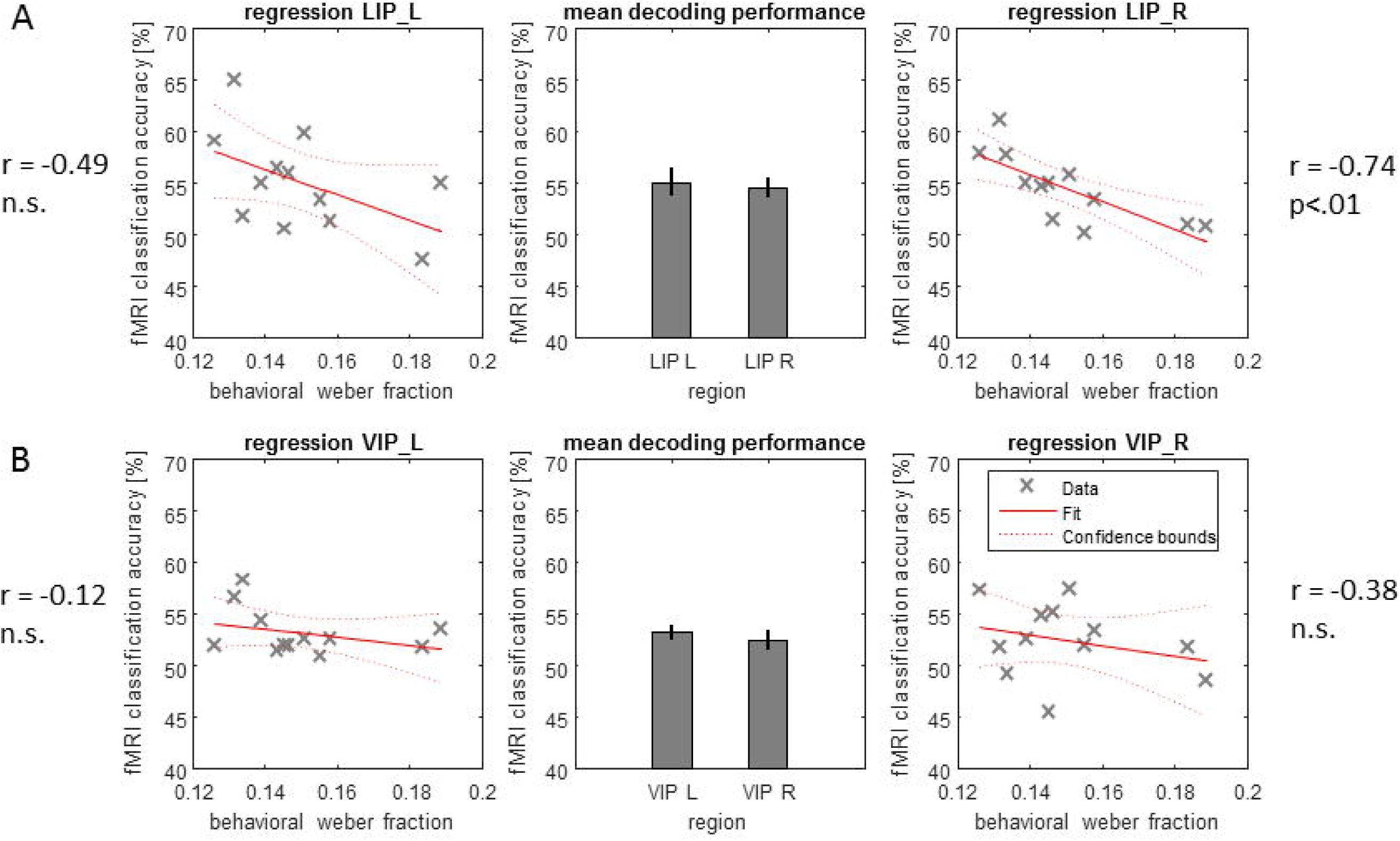
FMRI decoding results and across-subject correlation with behavioral Weber fractions in the left and right functional equivalents of areas LIP and VIP: Individual subjects’ ROIs were defined as the 150 voxels most activated across all sample numerosities vs baseline, within each one of the group ROIs as defined by a previous study (Eger et al., 2015). Panel A) displays the average classification performance (n=12, means and SEM) for pairwise discrimination between numerosities (theoretical chance level = 50 %) for left and right LIP (middle panel) together with correlations between individual subjects’ fMRI classification performance and behavioral Weber fractions (left panel for left hemisphere, right panel for right hemisphere). Panel B) displays the corresponding results for left and right VIP. The negative correlation between behavioral Weber fractions and fMRI decoding performance was only significant in the right LIP ROI when testing all regions in isolation.

In summary, correlation analyses revealed that subjects with better behavioral discrimination ability between individual numerosities also showed better decodability of numerosity evoked activity patterns. Critically, this tight relation was found in parietal cortex, and more specifically in the right LIP region, but not in occipital cortex, even though there was enough information in the latter area to decode numerosity with an equivalent level of performance.

### Behavioral precision and fMRI decoding performance across the numerical range

The results described previously allowed us to establish a relation between individual differences in behavioral numerical acuity and fMRI decoding performance, on average across all the numerosities tested. Behavioral results had also indicated some differences in behavioral precision across the numerical range (in particular smaller Weber fractions for 8 dots than for larger numerosities, see Figure 1D). In parietal cortex but not in occipital cortex these behavioral results were paralleled by a tendency for better decoding in the range including numerosity 8 (Figure 2A). To establish a more direct quantitative correspondence between these two tendencies, we computed the sensitivity index (d’) for all pairwise discriminations between numerosities from estimated behavioral Weber fractions on the one hand, and the output of the fMRI pattern classifier, on the other hand (see Materials and Methods for details). Figure 4 displays the resulting d’ matrices, for behavioral discrimination (A) and for decoding in early visual and parietal cortex ROIs (B), averaged across subjects. In spite of the fact that the overall discriminability of fMRI was much lower than the behavioral one, a general pattern of higher d’ for comparisons involving the lowest numerosity can be found in the behavioral matrix, and is more clearly reflected in the fMRI decoding matrix from the parietal than the early visual cortex ROI. Multiple regressions on the d’ values for the different numerical comparisons, run on a subject-by-subject basis (with predictors being the z-transformed ratio and mean size of the two numerosities in each pair, plus a constant) confirmed that for the behavioral data the beta weights for ratio and size were significantly different from 0 across the 12 subjects (paired two-tailed t-tests: ratio: t(11) = 23.81, p = 8.2e-11, size: t(11) = 3.16, p = 9.1e-03, Figure 3C). The same was true for decoding in the parietal cortex ROI (paired two-tailed t-tests: ratio: t(11) = 4.47, p = 0.0009, size: t(11) = 2.85, p < 0.0157), while for the early visual cortex ROI only the ratio predictor reached significance (t(11) = 5.21, p = 0.0003), but the effect of size remained non-significant (t(11) = 0.45, p = 0.6620) (Figure 4D).

**Figure 4:**
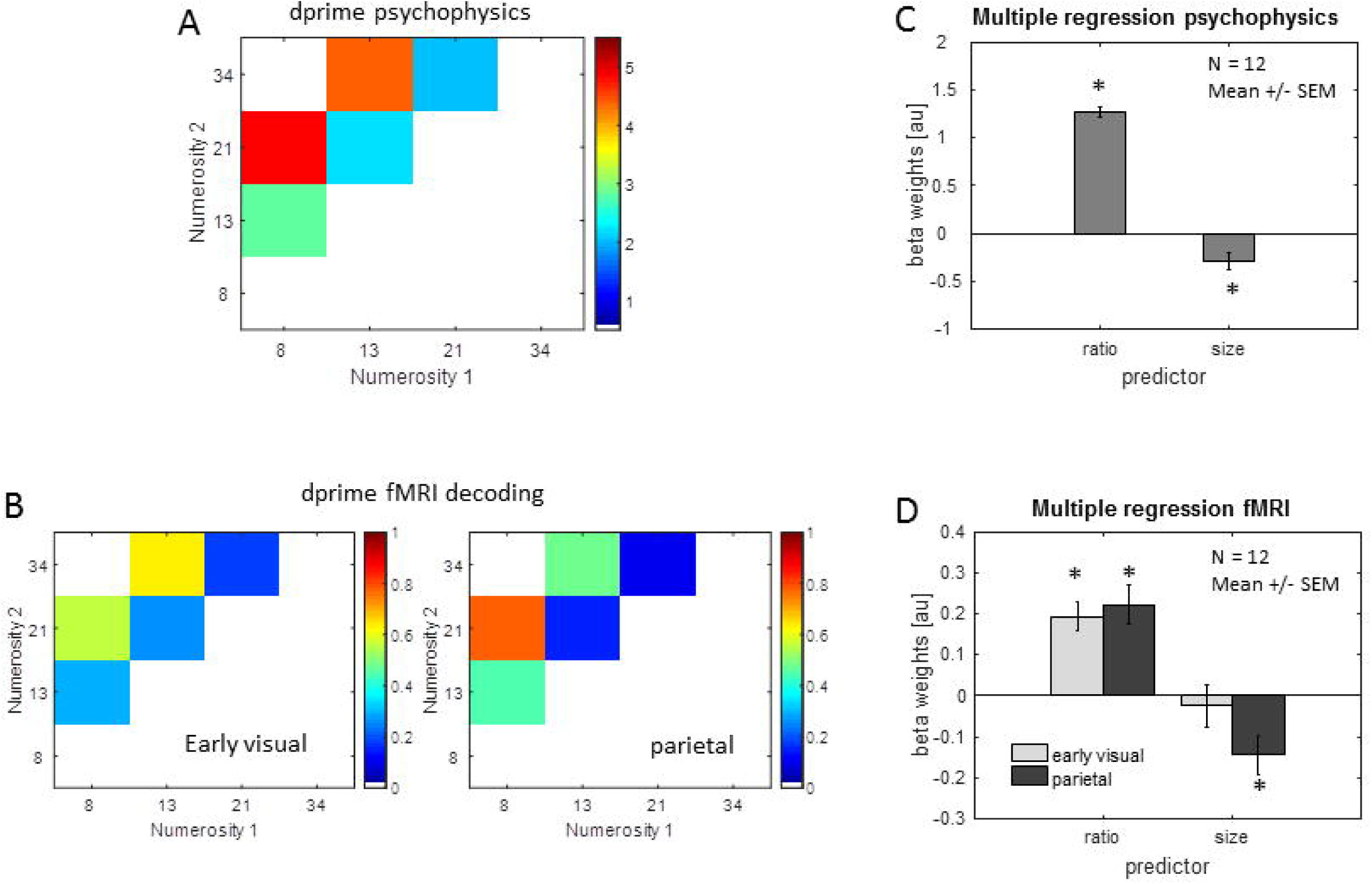
FMRI decoding results and behavioral precision across the numerical range: The d-prime index was calculated for all pairwise comparisons between numerosities (within each of the two numerical ranges) from behavioral Weber fractions and from fMRI decoding performance. Panel A) shows the across subject averages of the d-prime matrices for behavioral discrimination, and panel B) the ones for fMRI pattern discrimination in early visual and parietal ROIs. A tendency for higher d-prime values (indicating better discriminability) for comparisons involving smaller as opposed to larger numerosities can be seen in the behavioral results and in the fMRI results from parietal cortex. Multiple regressions, run on a subject-by-subject basis and tested for significance across subjects, confirmed a significant effect of numerical size on top of ratio, for the pairwise discriminability in the behavioral data (C) and the fMRI data of parietal cortex but not early visual cortex (D).

Thus, decoding performance in parietal cortex closely parallels the observed differences in behavioral discrimination performance across the numerical range, in particular showing a significant effect of the size of two compared numbers on top of their ratio, an effect absent in early visual cortex activity.

## Discussion

The present study acquired both psychophysical and functional neuroimaging data to test which of several regions previously shown to contain information about individual numerosities is read out when making explicit numerical judgements. We found that in spite of a very similar level of overall decoding performance in early visual and intraparietal cortex, the subjects’ behavioral Weber fractions measured prior to scanning were negatively correlated with pattern decoding performance in parietal (and more precisely in the right equivalent of LIP) but not occipital cortex. This result indicates that the precision with which subjects discriminate numbers reflects the precision with which they represent numbers in parietal cortex and hence demonstrates that this region is crucial for numerical decisions.

Overall, the fMRI decoding performance reported here (around 57 % correct in parietal cortex) may seem low compared to some previous studies (Eger et al., 2009; Bulthé et al., 2014; Knops et al., 2014). However, when considerably higher decoding performance was obtained previously this was generally for smaller numerosities (up to a maximum of 8 items). When the numerical values and ratios between numerosities were comparable to the ones used here, discrimination performance was considerably lower, and compatible with the one obtained in the current study (Eger et al., 2015). This also fits well with the fact that studies attempting to directly map the cortical layout of numerosity selectivity at the level of individual voxels have so far had sufficient sensitivity only to detect reliable responses for relatively small numerosities (Harvey et al., 2013; Harvey et al., 2017).

Given that early visual and parietal cortex allowed us in the present study to decode numerosity to the same degree, one might wonder what might be the nature of the information encoded in early visual cortex. We do not want to claim from the current results that early visual cortex does represent numerosity explicitely (implying individual neurons responsive or tuned to it at that level) and independently from other features. We used in this study two stimulus sets that either equated dot size or the total number of pixels across numerosities. While this is one commonly used strategy for stimulus creation in studies on numerosity perception, it does not control for differences in the total amount and distribution of contrast energy across numerosities, which can affect early visual responses. This factor may likely account for the early visual cortex results in this and other studies, given that the one previous fMRI decoding study which reported successful decoding of numerosity in parietal but not early visual cortex (Castaldi et al., 2016) had equated the contrast energy of the stimuli.

The LIP region defined here by neurophysiologically motivated localizers and showing a strongly significant relation with behavioral acuity is one of the parietal subregions that have been shown in the macaque monkey to contain neuronal responses discriminating between numerosities (Roitman et al., 2007), with VIP being another one where the majority of neurophysiological recordings were made (Nieder, 2016). Previously we have found that the human equivalents of both regions allowed for decoding of numerosity information for fMRI activity patterns (Eger et al., 2015). The fact that here we did not observe a significant relation of behavioral discrimination with VIP activation patterns does not necessarily mean that no such relation exists. Given the overall reduced decoding performance when the parietal ROI was replaced by several smaller subregions, null results can hardly be considered informative. Nevertheless, in the right LIP a significant relation was found in spite of the reduction in number of voxels, indicating that this region contains the most behaviorally relevant information at least at the spatial scale accessed by our measurements. It is interesting to note a close correspondence between the average MNI coordinates of our right LIP ROI (22.9 -61.0 54.2) and the coordinates (23 – 60 60) where a systematic spatial layout of numerosity responses was observed previously (Harvey et al., 2013) using ultra-high-field fMRI (though without testing the relation of these responses with behavioral numerical acuity).

The present report is the first one showing that the precision of intra-parietal (and more specifically right LIP) activity patterns evoked by individual numerosities reflects inter-individual differences in behavioral numerical acuity in adult human subjects. Interestingly, using fMRI adaptation methods, an across-subject correlation between behavioral Weber fractions and Weber fractions estimated from the fMRI adaptation effect was found very recently in 3-6 year old children (Kersey and Cantlon, 2017) in the right, but not the left intraparietal sulcus. FMRI adaptation and multi-variate decoding of evoked activity patterns are two different approaches often used to tackle similar questions (related to characteristics of neuronal representations), however, the underlying signals exploited by the two methods are of a very different nature. The fMRI adaptation approach is based on deviance signals which are observed at the level of regional activity when a change in the stimulus is introduced (Grill-Spector et al., 2006), possibly in line with an intrinsic tendency of the brain to predict its input (e.g., Kleinschmidt et al., 2002; Grotheer and Kovács, 2016). Multivariate decoding, on the other hand, exploits differences in direct evoked activation patterns, and therefore the layout of responsive neuronal populations, independently from temporal context/stimulation history. Given that such different measures cannot necessarily be guaranteed to provide the same insights, and that they can in fact sometimes lead to different results in the same paradigm and data set (Drucker and Aguirre, 2009), it is even more remarkable that in the case of the relation between perceptual and neuronal sensitivity to numerosity studied here, both measures converge across age groups onto similarly located parietal regions. Beyond this work using fMRI, it is of interest that an across subject correlation has also been observed between the amplitude of ERP components (in particular the N2pc) and numerosity discrimination within the subitizing range, however without allowing to localize the underlying brain regions generating this effect (Ester et al., 2012).

While as far as we are aware of, inter-individual differences in numerical acuity have not been investigated at the level of single neuron responses, macaque neurophysiology has nevertheless provided evidence for the behavioral relevance of the intraparietal and prefrontal number selectivities: specifically, on trials where the monkeys made errors in a delayed numerosity match-to-sample task, the numerical selectivity for the preceding sample stimulus was lost or reduced (Nieder and Miller, 2004). A related approach using error trials to reveal the relevance of regional activation patterns for performance has also been used in fMRI studies related to other cognitive domains than number: for example, in an experiment on object recognition, on trials when subjects failed to correctly identify the object that was briefly presented and masked, discriminative information was disrupted in lateral occipital object responsive areas, but not in early visual cortex (Williams et al., 2008). Such effects are not necessarily restricted to higher-level cortical areas: decoding of stimulus orientation from early visual cortex was shown to be enhanced on trials where subjects correctly discriminated a small change in orientation as compared to incorrect trials (Scolari and Serences, 2010), and the uncertainty about orientation decoded from early visual cortex on individual trials was found to reflect the variability of perceptual decisions (van Bergen et al., 2015).

The paradigm we used was less suited to investigate the effects of behavioral relevance at the level of individual trials: subjects were performing a delayed numerosity comparison with two response alternatives (larger or smaller) for which chance performance is 50 % and furthermore errors could either arise at the level of encoding of the first or second stimulus, thus minimizing the chances to identify true errors with the amount of trials available in an fMRI experiment as this one. We therefore focused on differences in behavioral precision across subjects and their relation with fMRI decoding. Only one other study to our knowledge has so far reported that decoding performance for a fine-grained perceptual comparison predicted inter-individual differences in behavioral discrimination capacity: in a phoneme discrimination task, decoding of the phonemes /ra and /la differed between English and Japanese speakers, but in addition was also indicative of individual differences in behavioral acuity within groups (Raizada et al., 2010). Interestingly, and different from our results, behavioral acuity in that study was related to representations at the earliest stage of cortical processing (Heschl’s gyrus).

The across-subject relation with behavioral performance in our case was found for numerosity, but since only numerosity was tested, we cannot claim that the relation is specific for that feature in the regions in question, rather than potentially more generally observed for relevant contents in a comparison task. However, establishing the specificity of the relation between behavior and fMRI decoding for numerosity per se appears a non-trivial enterprise, since successful decoding of numerosity already requires a considerable amount of data/scanning time, and any additional feature to be contrasted with numerosity regarding its correlation with behavioral performance would need to be discriminable within the same regions in the first place.

A related issue is whether the correlation observed necessarily needs to reflect the subjects’ intrinsic numerical acuity, rather than variations across subjects in the level of engagement in, or attention to, the task. Attention is well-known to modulate activity in parietal cortex, and pattern recognition methods have revealed that distributed response patterns in intraparietal areas not only can reflect task set (which feature dimension is attended), but also preferentially discriminate between feature values within an attended/task-relevant dimension (e.g., Liu et al., 2011; Ester et al., 2016). Nevertheless, such attentional enhancement of feature information has been found to be at least as present at the earliest stages of the visual hierarchy (e.g., Serences and Boynton, 2007; Jehee et al., 2011; Ling et al., 2015). This contrasts with our study, where we find that only parietal and not early visual cortex patterns reflect behavioral acuity for numerosity across subjects, although our design does not allow us to explicitly rule out effects related to attention in this finding.

In addition to the across subject relation between behavioral acuity and pattern decoding performance for numerosity discrimination, our study also observed differences in numerical acuity across the numerical range tested. Specifically, once again in parietal, but not early visual cortex, smaller numerosities were better discriminated than larger numerosities of the same ratio, paralleling better behavioral precision for the smaller numerosites. Similar findings showing that small numerosities are more discriminable than predicted by Weber’s law have been obtained in a few other behavioral studies (Merten and Nieder, 2009; Burr et al., 2010). Interestingly, Burr et al. (2010) have shown that the higher behavioral precision for smaller numerosities disappeared when attentional resources were engaged elsewhere, leading to the suggestion that several potential mechanisms contribute to numerosity discrimination across the numerical range, one of which (for relatively small numerosities) requires attentional resources. In our study, the better discriminability of smaller numerosities was not restricted to numbers as small as the ones used in that study and it still remains to be understood in detail which task or stimulus factors determine up to which limit performance for smaller numerosities can deviate from Weber’s law.

The findings reported here were obtained during delayed number comparison and thus a working memory task. Further studies will be needed to clarify whether the present correlations between behavioral numerical acuity and the precision of the neuronal representation reflects the precision of the numerical percept per se, or of its maintenance in short-term memory. On the other hand, a previous study has lent support to the idea that the mechanisms for extracting numerosity and non-numerical feature tracking/short term memory of properties of visual sets might be closely intertwined (Knops et al., 2014): Area LIP in particular is thought to implement a saliency map, and a computational model of a saliency map architecture did account for fMRI data acquired with two different tasks in that region by different levels of mutual inhibition between model nodes. It seems tempting to speculate how such a shared component could potentially provide a unifying explanation for diverse impairments found in disorders of numerical processing (dyscalculia) that are conventionally categorized as either domain-specific (impairments in non-symbolic number processing) or domain-general (impairments in visual working memory) (Piazza et al., 2010; Szucs et al., 2013).To summarize and conclude, the present study is the first one to demonstrate that the precision of numerosity evoked activity patterns in intraparietal cortex, and more specifically the equivalent of area LIP, correlates with behavioral enumeration abilities and thus likely constitutes a crucial level of the cortical hierarchy at which activity is read out for perceptual decisions during numerosity tasks. Future studies may clarify whether this link between behavioral precision and evoked response patterns in intra-parietal cortex is specifically found for numerosity or also for other quantitative or even non-quantitative perceptual contents, what is the role of attention to or task relevance of the feature in question, and what, if any, is the relation between the brain behavior correlation described here and higher-level aspects of numerical performance.

## Footnotes

1 We also tested in how far Weber fractions differed as a function of the stimulus set (dot size vs total surface area equated between numerosities). In an analysis performing separate fits per stimulus set, magnitude and range, an ANOVA revealed a main effect of stimulus set (F(1,11)=17.2, p=0.0016, with average w being 0.155 for the constant dot size set, and 0.172 for the constant total surface area set) which, however, did not significantly interact with the other factors. Investigating in detail the influence of stimulus properties on numerical discrimination is beyond the scope of the present study, and for reasons of sensitivity / increasing the number of trials, we subsequently collapsed across the two stimulus sets in both the behavioral and fMRI analyses.

